# Research on the Influence of Different Temperature Conditions on the Microbial Community in Anearobic-Anoxic-Oxic process on the Plateau

**DOI:** 10.1101/2020.09.01.277244

**Authors:** Decai Huang, Yongchen Zong, Ning Zhang, Yuanwei Li, Kaiyue Hao

## Abstract

In order to further understand the influence of different temperature conditions in the low temperature range on the microbial community structure in the Anearobic-Anoxic-Oxic waste water treatment process on the plateau, four temperature conditions were designed in the research, including 25°C, 20°C, 15°C and 10°C. Each working condition lasted three days. Activated sludge from anaerobic tank, anoxic tank and oxic tank under each working condition was analyzed based on the 16S rRNA high-throughput sequencing technology. The result shows that the plateau temperature conditions have little influence on the level abundance of phylum. Under all conditions, Bacteroidetes, Proteobacteria and Actinobacteria are the main phyla. The abundance of nitrifying bacterium and phosphorus removal bacteria rose in the first three conditions and fell at 10 °C. The abundance of Denitrification bacteria and Nitrite oxidizing bacteria was significantly higher than that of Ammonia oxidation bacteria and Azotobacter bacteria and Phosphorus-accumulating Organisms(PAO) had an overall advantage over Glycogen-accumulating Organisms (GAO) throughout the research.

## Introduction

86.1% of Tibetan area is located at an elevation of more than 4000mabove sea level**[1]**, and the high-altitude areas feature unique climatic conditions. According to the bureau of meteorology, the average daily temperature of Tibetan area is between 3°C and 17°C, and outdoor temperature varies widely from day to night. The average daily temperature difference exceeds 10°C. The perennial low temperature and the temperature difference from day to night in Tibet are the climatic phenomena different from those of other regions, which make the rules of waste water treatment and the microbial community structure of activated sludge on the plateau more complex than those of other regions **[2]**.

According to the relevant metabolic functions involved in waste water treatment, the microbial communities in activated sludge are divided into ammonia oxidation related functional bacteria genera, nitrite oxidation related functional bacteria genera, denitrification related functional bacteria genera, biological phosphorus removal related functional microbial genera, methane metabolism related functional microbial genera and heterotrophic metabolism related functional microbial genera **[3]**. The removal of nitrogen and phosphorus from waste water treatment is generally studied. According to existing studies, temperature has a complex influence mechanism and effect on nitrogen removal related functional microbial genera, such as Ammonia-oxidizing bacteria, Commanmox, Ammonia-oxidizing archaea, Nitrite oxidizing bacteria, Denitrifying bacteria, and biological phosphorus removal related functional microbial genera such as polyphosphorous bacteria and glycan bacteria **[4]**.

Because functional bacteria are sensitive to the changes of temperature, temperature has a great influence on the effect of biological waste water treatment,such as Anearobic-Anoxic-Oxic(A2O) process. According to the research of Li, C[5], the effect of temperature on the removal rate of COD(Chemical Oxygen Demand) and TP(Total Phosphorus) is not obvious. In the low temperature stage, a high and stable removal rate can be achieved by extending the sludge age or adding carbon source. The temperature rise within the appropriate range will significantly increase the removal rate of TN(Total Nitrogen) and NH4+-N, according to the study of ZHANG.X.D **[6]** When the temperature rises from 8°C to 26°C, the removal rate of TN and NH4+-N increases from 16%, 18% to 84% and 69% respectively, and the optimal temperature is between 25°C and 30°C.

Studying the removal effect alone does not reveal a deeper mechanism. Based on the fact that most bacteria cannot be purified alone **[7]**, the metabolic reactions of related bacteria are complex, and the variability of bacteria itself is strong, it is prone to draw wrong conclusions from the study of microbial community changes. By combining the removal effect with the change of microbial communities, the influence of environmental changes on the microbial communities of activated sludge can be more clearly revealed.

## 1. Materials and methods

### 1.1 Experimental equipment and methods

With temperature controlled, A2O waste water treatment technology on plateau was adopted to study the influence of special temperature conditions on the microbial community of A2O process in the plateau. Four A2O waste water treatment equipment are mainly made of acrylic. The volume ratio of anaerobic tank, anoxic tank and oxic tank is 1:1:2 and the total volume is 40 L. The waste water treatment experiments were carried out under the conditions of 25°C, 20°C, 15°C and 10°C to simulate the temperature conditions on plateau and each working condition lasted for 3 days. With the use of temperature control switches to control the heating rods, the temperature condition was controlled. The DO (Dissolved Oxygen) of Oxic tank was controlled at 3.0±0.2mg/L with the use of air pump and gas flow meter. The mixture in anoxic pool was refluxed through a peristaltic pump with a reflux ratio (Ri) of 200%. Sludge reflux was controlled by magnetic circulating pump, and sludge reflux ratio is R=100%. Samples were taken from the influent, effluent, anearobic tank, anoxic tank and oxic tank every 8 hours for water quality testing. Sludge samples were taken from anearobic tank, anoxic tank and oxic tank after water samples were taken for the last time in each working condition.

### 1.2 Influent in the experiment

Waste water from a septic tank on a campus in Tibet was used for the experiment. The water intake point and the location of the experiment were 3000m above sea level, and the air pressure was 0.71*101.325 kPa (value of standard atmospheric pressure). Temperature of the influent was between 8.1 and 14.7 °C. DO was between 0.09 and 0.16 mg/L. And pH was between 6.93 and 7.62. As for the four chemical data, COD was between 217 and 526 mg/L, TN was between 65.1 and 112.5 mg/L, TP was between 2.96 and 6.81 mg/L, and NH_4_^+^-N was between 2.96 and 6.81 mg/L. The influent is a typical low C/N ratio waste water **[8]**.

### 1.3 Testing methods

The temperature, pH and DO of waste water were detected with the water quality quick meter (HQ40d). COD, TN(Total Nitrogen), TP and NH4+-N were detected with water quality analyzer (model: 6B-3000A) produced by Jiangsu Shengaohua Company. COD was measured by Potassium Dichromate method**[9].** TN was measured by Potassium persulfate digestion-Ultraviolet Spectrophotometry **[10].**TP was measured by Ultraviolet Spectrophotometry **[11].** And NH_4_^+^-N was measured by Napierian Reagent Colorimetric **[12].**

### 1.4 Biological detection methods

The biological data were sequenced by 16S rRNA gene by Shanghai Majorbio Bio-pharm Technology Co.,Ltd. In each working condition, the anaerobes tank, anoxic tank and oxic tank were sampled once at the end of each working condition. 50ml mixture of mud and waste water in each tank was taken and put into the centrifuge tube and centrifuged at 5000 RPM for 15 minutes in a small centrifuge. The 30mL supernatant was removed. After the centrifuge tube containing the layered 20mL mixture was sealed with sealing film, the centrifuge tube was frozen at −80°C for 12 hours, then the sample was sent to the company by post. The v3-v4 variable region was amplified by PCR using a biological DNA extraction box E.Z.N.A.^®^ Soil DNA Kit. The primers used were 515F(GTGCCAGCMGCCGCGGTAA) and 907R(CCGTCAATTCMTTTRAGTTT). The amplified samples were sequenced on the Illumina Miseq sequencer.

## 2. Results and analysis

### 2.1 Removal rate of carbon, nitrogen and phosphorus

As shown in figure 1, Origin 2016 was used to plot the removal rates of COD, TN, TP and NH4^+^-N of effluent from the continuous experimental data of four working conditions. Each point stands for one chemical data of The COD removal rate of effluent was between 69.34% and 92.46%. The highest removal rate was 92.46% in the working condition of 15**°C**. and the lowest was 70.48% at 20°C. When the temperature was adjusted from 25°C to 20°C the removal rate decreased by 14.85%, which was the largest decrease. After the domestication in the working condition of 20°C and 15°C, the removal rate gradually stabilized. In the working condition of 25°C the average removal rate of a single working condition was highest, reaching 83.59%. And in the working condition of 20°C, it was 73.13%, which was the lowest. The average absolute deviation can reflect the stability of the data which reflects the stability of the working condition **[13]**. The smaller the value is, the stabler the working condition is. According to that, the lowest value occurred at 10°C, which was 1.73, and the highest value occurred at 15°C, which was 4.510. The overall dispersion degree of COD removal rate was 6.03.

**Figure 1.**
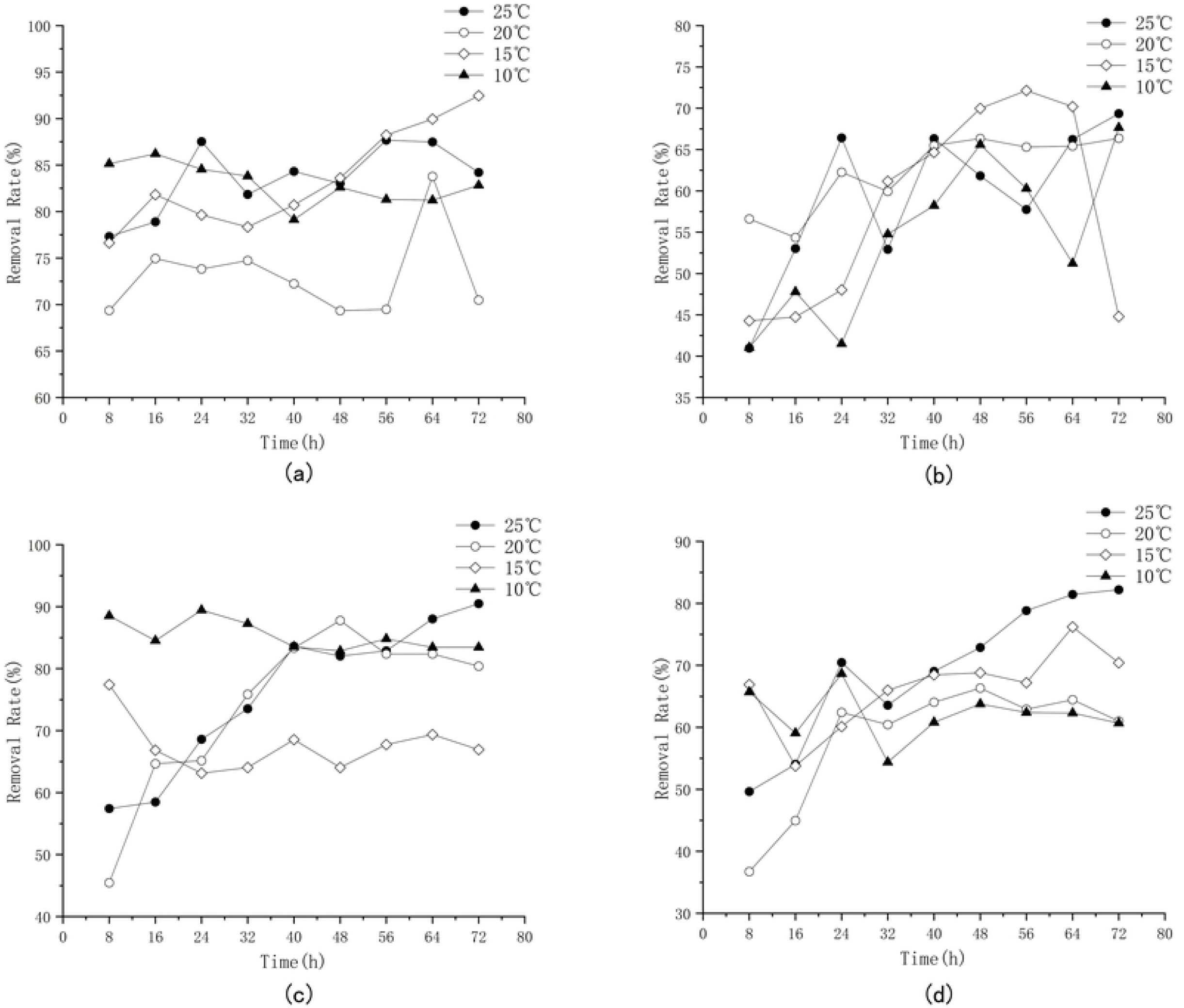
Pollutants removal rate during the continuous operational period

The TN removal rate of the effluent was between 40.97 and 72.14%, with the highest value at 20°C (62.45%) and the lowest value at 10°C (54.23%). The lower the temperature was, the longer the recovery time of removal rate would be, and the lower the maximum removal rate would be in the working condition. The average removal rate of single working condition was highest at 20°C (62.46%) and lowest at 10°C (54.23%). The lowest value of dispersion was 4.51 at 20°C, and the highest value was 12.16 at 15 **°C**. The overall dispersion degree of TN removal rate is 9.39.

The TP removal rate of the effluent was between 45.45% and 90.50%, the highest value was 90.50% at 25°C, and the lowest value was 45.45% at 20°C. When the TP removal rate was adjusted from 25°C to 20°C, it decreased significantly, but at 10°C, it showed a stable high removal rate. The average removal rate of a single working condition was highest at 10°C (85.31%) and lowest at 15°C (67.56%). The lowest value of dispersion was 4.51 at 20°C. and the highest value was 12.16 at 15**°C**. The overall dispersion degree of TP removal rate was 11.08.

The NH4+-N removal rate of effluent was between 36.73% and 82.21%, with the highest value at 25°C, 82.21%, and the lowest value at 20°C, 36.73%. NH4+-N removal rate is less affected by temperature change than that of TN. The average removal rate of single working condition was highest at 25°C (69.12%) and lowest at 20°C (58.14%). The lowest value of dispersion was 4.05 at 10°C, and the highest value was 11.57 at 25°C. The overall dispersion degree of NH4+-N removal rate was 9.24.

Consistent with the research results of **[14]**, the removal rates of COD and TP are less affected by temperature, while the removal rates of TN and NH4+-N are more affected by temperature. The overall removal rate of COD is relatively stable. The removal rate of TP is sensitive to temperature changes above 15°C, and it fluctuates greatly. With the decrease of temperature, the stability gradually improves. Removal rates of TN and NH4+-N are greatly affected by temperature. The removal rate of TN and NH4+-N decreased significantly when the temperature was lower than 15°C.

### 2.2 Waste water treatment index discharge grade

In table one, the value outside the brackets is the control index when the water temperature is above 12 °C, and the value inside the brackets is the control index when the water temperature is below 12 °C. The total phosphorus standard value takes the sewage treatment plant built since January 1, 2006 as the standard.

According to the national standards of the People’s Republic of China (GB18918-2002) **[15]** for the discharge of pollutants from urban waste water treatment plants, the discharge grades of waste water treatment indicators in the whole working condition were included in a table, as shown in Table 1. The COD value of effluent reached the highest standard rate of primary standard, which was 66.75%. The TP value of effluent reached a lower standard rate of 41.67%. The TN value and Nh4+-N value reached the lowest in the first class standard, which were 11.1% and 0% respectively. The temperature conditions at the plateau are unfavorable for nitrogen removal. On the whole, the effluent from waste water treatment reached the highest standard rate at 20°C.

**Table 1.** The standard times the maximum allowable emission concentration of basic control project reaches

| Basic control item |  | Primary standard |  | Secondary standard | Tertiary standard |
| --- | --- | --- | --- | --- | --- |
|  |  | A standard | B standard |  |  |
| COD | Standard value (mg/l) | 50 | 60 | 100 | 120 |
|  | Times reach the standard | 17 | 7 | 10 | 1 |
| TN | Standard value (mg/l) | 15 | 20 | — |  |
|  | Times reach the standard | 2 | 2 | 32 |  |
| NH<sub>4</sub><sup>+</sup>-N<sup>①</sup> | Standard value (mg/l) | 5 (8) | 8 (15) | 25 (30) | — |
|  | Times reach the standard | 0 | 0 | 14 | 22 |
| TP<sup>②</sup> | Standard value (mg/l) | 0.5 | 1 | 3 | 5 |
|  | Times reach the standard | 3 | 12 | 15 | 1 |

### 2.3 Overall change of microbial diversity

#### 2.3.1 Diversity analysis of activated sludge samples

16S rRNA high-throughput sequencing technology was used to distinguish the data of each sample through the Index sequence, and fastp and FLASH software were used to control and filter the quality of reads and the effect of merge, so as to optimize the data. A total of 522,177 high quality sequences were obtained from activated sludge samples for subsequent analysis under 4 temperature conditions, with the length of high quality sequences distributed in 401-440nt, mainly concentrated in 420-440nt. In contrast with Silva databases, Usearch software was used to carry out a repetitive sequence (not including single sequence) OTU cluster according to 97% similarity while removing chimera in the process of clustering. The QIIME software was used to count the microbial information of activated sludge according to the taxonomy(Table 2), and calculate the OTU and genus of the four temperature conditions (Figure 2 **a&b**).

**Table 2.** Taxonomic statistics of activated sludge samples

| Conditions | Phylum | Class | Order | Family | Genus | Species | OTU |
| --- | --- | --- | --- | --- | --- | --- | --- |
| 25°C | 28 | 49 | 120 | 228 | 477 | 694 | 962 |
| 20°C | 29 | 58 | 138 | 256 | 553 | 858 | 1191 |
| 15°C | 29 | 53 | 129 | 249 | 541 | 826 | 1139 |
| 10°C | 29 | 49 | 124 | 232 | 485 | 724 | 990 |

**Figure 2.**
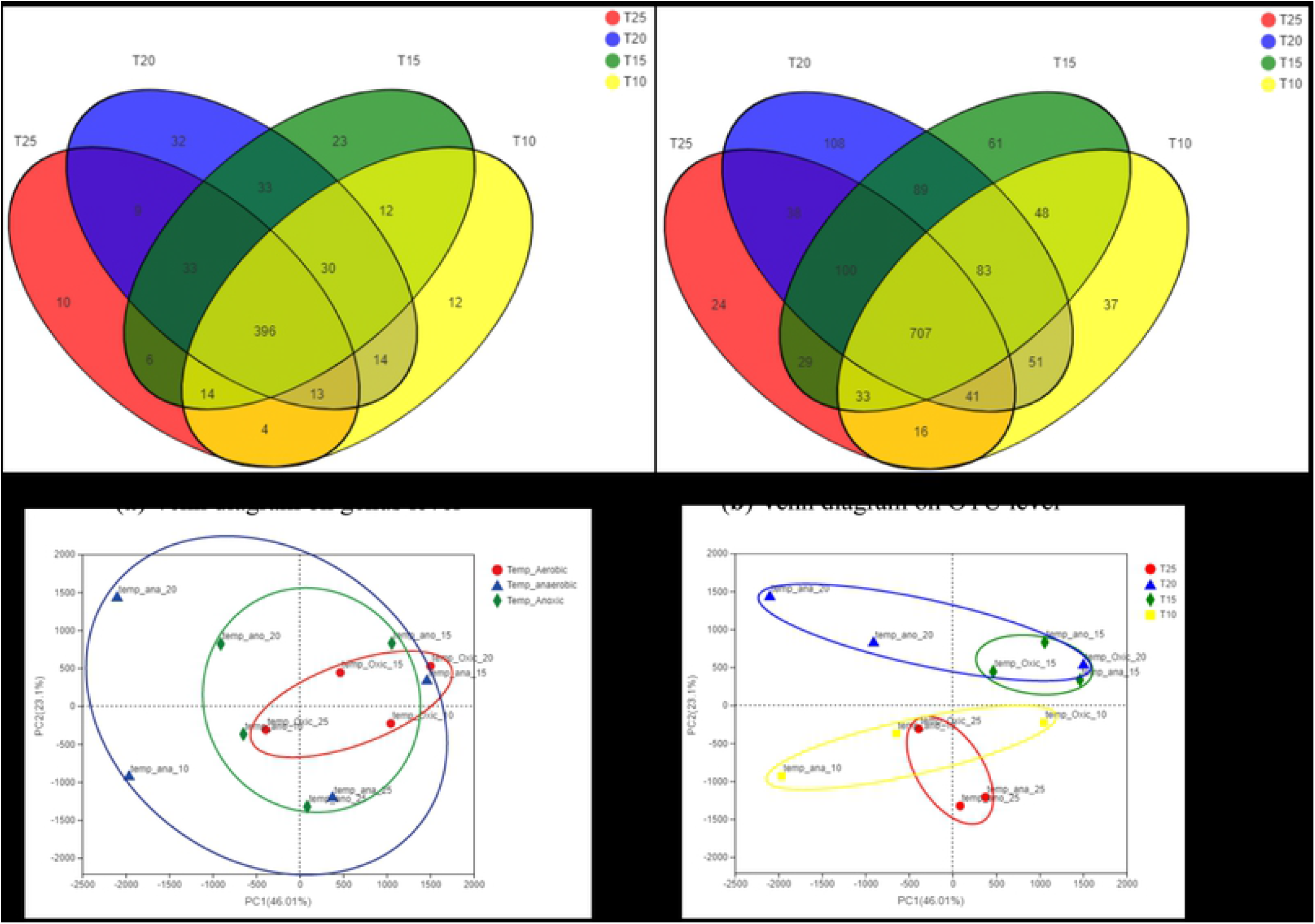
Difference analysis between groups

As can be seen from Table 2, the OTU numbers at 20°C and 15°C are 1191 and 1139, respectively. The OTU numbers at 25°C and 10°C are 962 and 990 respectively. The species richness at 20°C and 15°C was significantly greater than that at 25°C and 10°C. At 25°C, the effluent stability was the worst, which was directly related to the low richness of the microbial system. In the Venn diagram on OTU level and genus level, it can be seen that among the 1445 OTUs detected, 707 are detected in all conditions, and 122 are only detected in one condition. Among the 636 Genus detected, 396 are detected in all conditions, and 77 are only detected in one condition. Although the temperature conditions change greatly and the treatment effect varies greatly, the microbial community of activated sludge was stable on the whole, which could maintain the basic function of denitrification and dephosphorization. PCA, or principal component analysis, is used for dimensionality reduction analysis of data correlation,which can reflect the correlation between the data**[16]**. The analysis of grouping by pool and the analysis of grouping by working condition were carried out(Figure 2 **c&d**). Obviously, the data aggregation of the microbial communities grouped according to the working condition was better, and the influence of temperature on the microbial communities was greater than that caused by the different environments of different ponds. So, in the subsequent discussion, the sample groups under different working conditions were discussed as a whole as far as possible.

The sequencing coverage, richness and diversity of activated sludge under different temperature conditions were analyzed through α diversity analysis. Coverage index(Coverage), richness index (Sobs) and diversity index (Simpson, Shannon) of species sequencing were taken, and the results were shown in Table 3.

**Table 3.**
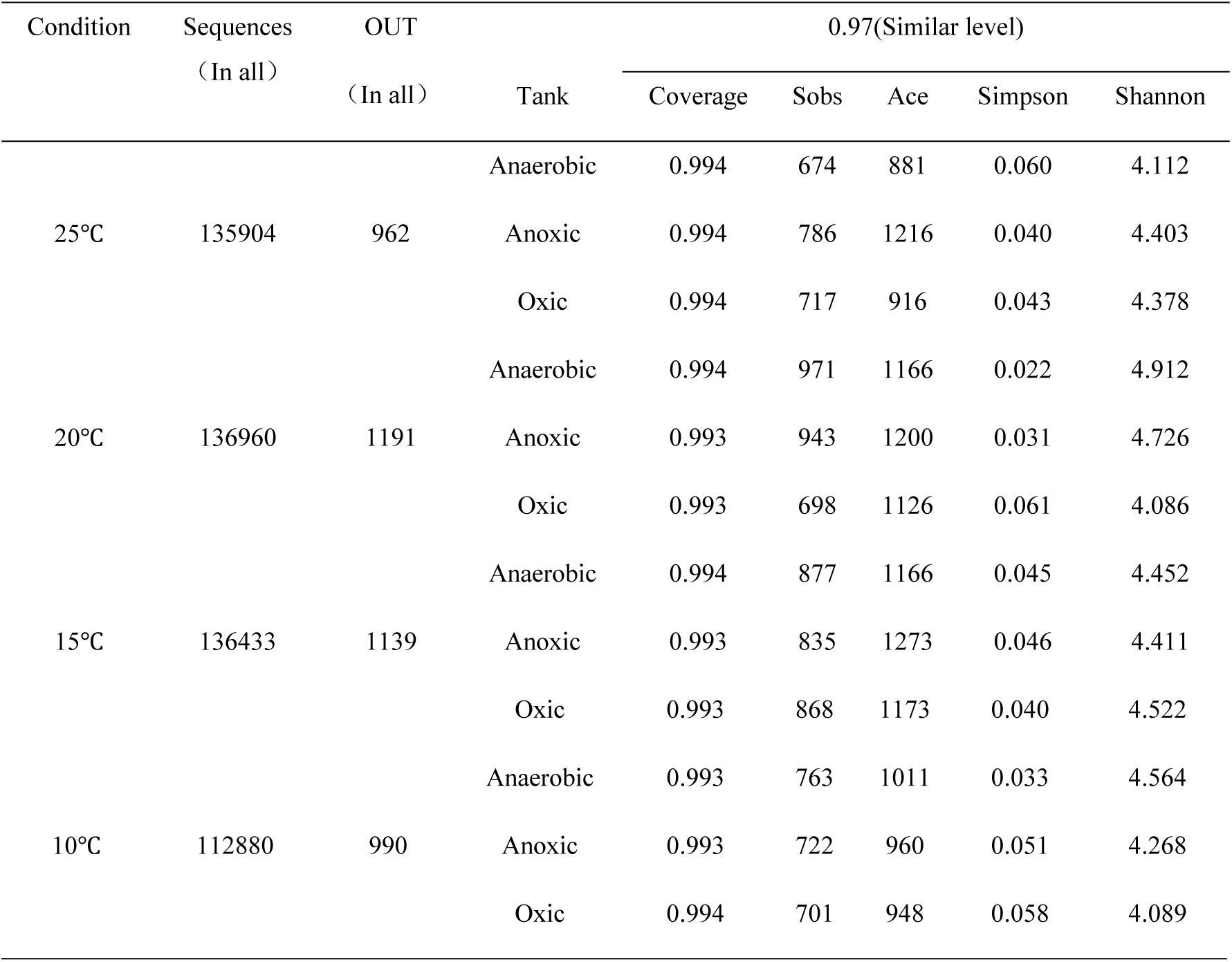
The α diversity analysis of activated sludge samples

The Coverage indexes of 12 samples in 4 working conditions were all greater than 0.993 **[17]**, which could effectively represent the information of microbial communities in this experiment. Sobs index reflects the actually observed value of richness **[18]**. Combining the OTU number, the richness at 25°C and 10°C is worse than that at 20°C and 15°C. By comparing the samples under various working conditions, it can be seen that the Ace index and Shannon index at 20°C and 15°C were higher than those at 25°C and 10°C, while the Simpson index at 20°C and 15°C were significantly lower. The Ace index and Simpson index are mainly used to measure the diversity of samples and reflect the microbial richness in samples. As the Ace index gets smaller, the Simpson index gets greater, and species diversity in the system gets lower. In addition to assessing microbial diversity in the sludge, Shannon index can also be used to evaluate its uniformity **[19]**. It can be seen that the homogeneity of microorganisms at 25°C and 10°C was low in this experiment. At 25°C and 10°C, the diversity, richness and homogeneity in the activated sludge system were all lower. So it can be speculated that under these two temperature conditions, the development of some strains is obviously better than that of others. When the temperature was 25°C, the poor stability of the waste water treatment effect was related to the obvious advantages of some strains. At 10°C, the removal capacity of nitrogen decreased significantly, and the number of species, diversity, richness and uniformity of activated sludge decreased, which may be related to the decrease in the number and abundance of bacteria related to the removal function of nitrogen.

#### 2.3.2 Variation of the diversity of microbial communities on phylum level

According to the hydraulic residence time ratio of 1:1:2, the microbial community composition in the anaerobic tank, anoxic tank and oxic tank was combined according to the ratio of 1:1:2. The microbial community composition on phylum level under the condition of plateau temperature was shown in Figure 3, and the phyla with less than 1% abundance were incorporated into Others. In all the four temperature conditions, more than 20 phyla were incorporated into Others, and the total proportion was less than 1.5%. The dominant bacteria on phylum level are very obvious. The top three are Bacteroidetes (31.8%-33.9%), Proteobacteria (27.5%-33.4%) and Actinobacteria (17.9%-23.6%). Proteobacteria, Bacteroidetes and Actinobacteria are the dominant bacteria in many waste water treatment and natural waste water researches. It has been reported that they have strong adaptability to the environment **[20–23]**, like the strong adaptability to growth inhibiting environments such as excessively high and low temperature, antibiotics and high COD. And to explore the influence of rapid temperature change, the single working condition lasts a short time, so it is reasonable that the overall change is small. Other dominant bacteria Firmicutes, Choloroflexi, Patescibacteria, Acidobacteria, Verrucomicrobia have been reported in other studies on activated sludge **[24]**. Accounting for more than 85%, Bacteroidia from Bacteroidetes, Clostridia from Bacteroidetes, Actinobacteria from Actinobacteria, α-Proteobacteria from Proteobacteria, β-Proteobacteria and γ-Proteobacteria are the dominant bacteria on class level. Theδ-Proteobacteria class from Proteobacteria phylum, which is commonly reported together with α-Proteobacteria and β-Proteobacteria, was less than 1.5%. Theδ-Proteobacteria accounted for nearly 1.4% at 15°C and less than 0.8% in other conditions. With the decrease of temperature, the change rules of dominant class were not consistent. For example, Bacteroidia and Actinobacteria decreased first and then increased, β-Proteobacteria and α-Proteobacteria rose first and then decreased, and β-proteobacteria and α-Proteobacteria decreased at 15°C and 10°C respectively. However, the total proportion of the six dominant classes was 85±5%, showing a certain stability.

**Figure 3.**
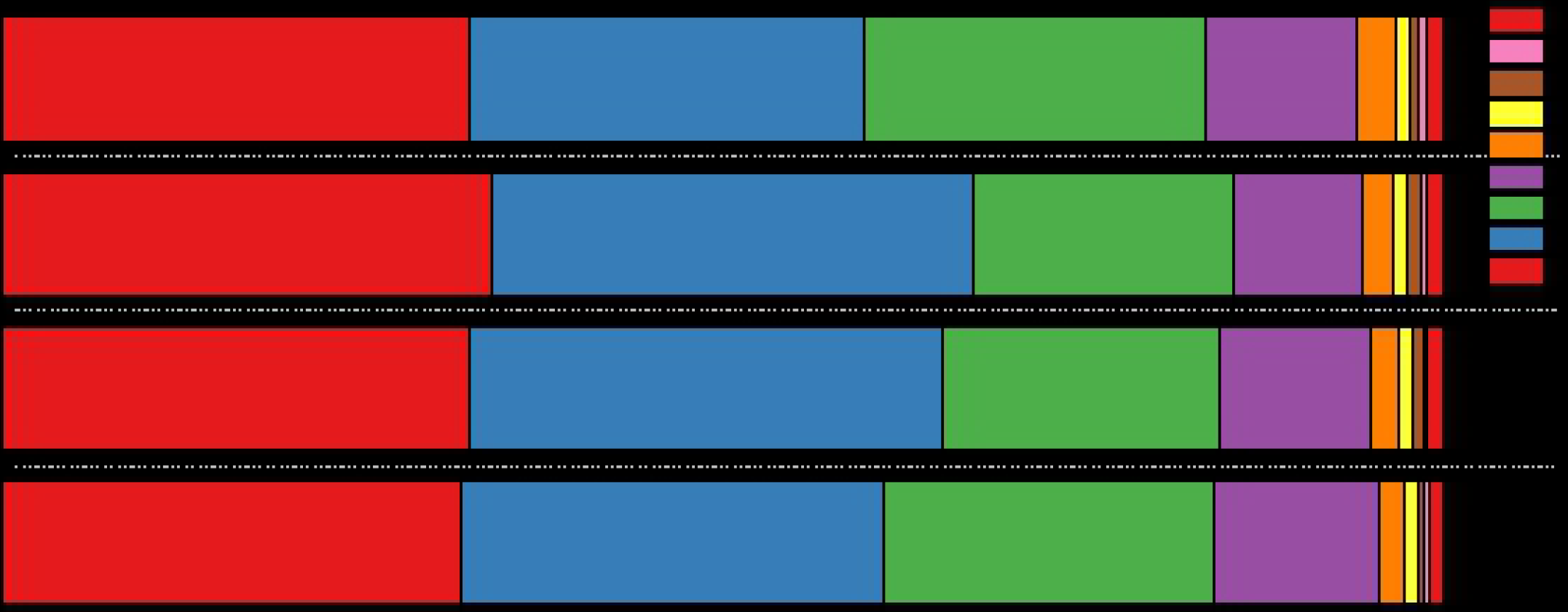
Bar chart of microbial community on phylum level

#### 2.3.3 Changes in the overall abundance of denitrifying and dephosphorizing bacteria

In order to further understand the effect of rapid temperature change conditions on the functional bacteria community of denitrification and phosphorus removal in activated sludge. The 100 genera with the highest horizontal abundance were counted to draw the Heatmap diagram. The similarity and difference of the community composition of activated sludge samples under different temperatures on genus level were reflected through color ladder and similarity degree, as shown in Figure 4. The genus was marked on the left side of each row of Heatmap for similarity cluster analysis, and the Denitrification bacteria related to nitrogen removal and Denitrifing phosphorus removal bacteria related to phosphorus removal were marked with blue frame and red frame.

**Figure 4.**
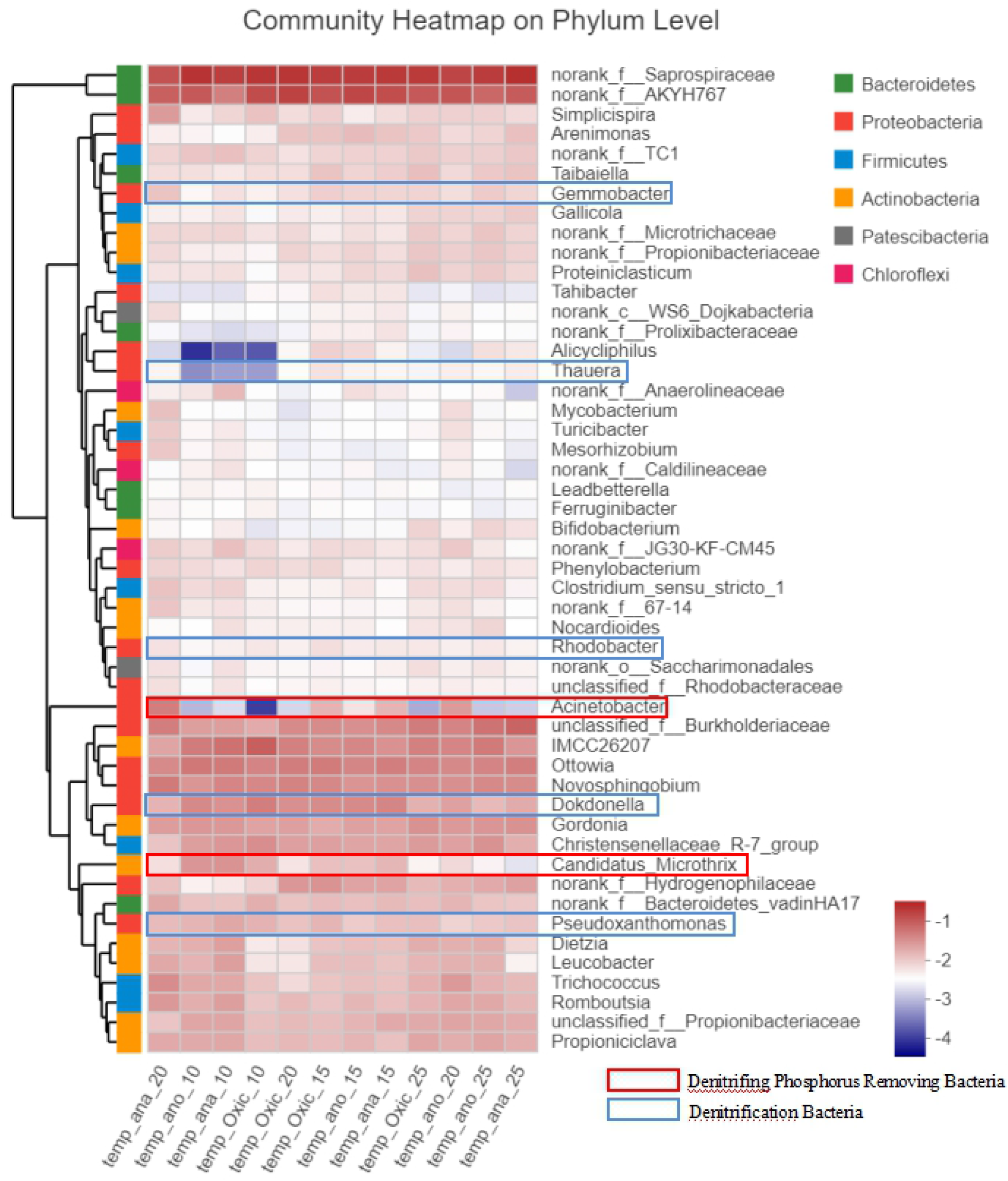
Heatmap diagram on genus level

In the first 100 genera of abundance, there were 5 genera related to the removal function of nitrogen, including Pseudoxanthomonas, Gemmobacter, Thauera, Rhodobacter, Dokdonella, and Pseudoxanthomonas, and 2 genera related to the removal function of phosphorus, including Acinetobacter and Candidatus_Microthrix. Five dominant genera were all from Proteobacteria, mainly from α-Proteobacteria, β-Proteobacteria and γ-Proteobacteria and a total of 16 related OTUs were detected in this sequencing. Rhodobacter from γ-Proteobacteria was found to have the largest OTU number (5). The two dominant genera related to the removal function of phosphorus were all from γ-Proteobacteria from Proteobacteria. In this sequencing, 10 related OTUs were found, of which 7 were Acinetobacter. As can be seen from Figure 4, the genera related to phosphorus removal function are relatively concentrated, while the bacteria related to nitrogen removal function are relatively dispersed. The genera related to the removal function of phosphorus are mainly divided into Phosphorus-accumulating bacteria (PAO) and Glycogen-accumulating bacteria(GAO) and the function is relatively simple while the genera related to the removal function of nitrogen removal can be divided into Ammonia oxidizing bacteria(AOB), Commamox, Ammonia-oxidizing archaea(AOA), Nitrite oxidizing bacteria(NOB), Denitrifying bacteria(DNB), and their functions and reactions are much more complex than those of phosphorus removal bacteria **[25]**.

According to the role played in nitrogen removal function, the Denitrification bacteria found in this sequencing(Figure 5 **a**) could be divided into 4 genera, Azotobacter **[26]**, AOB **[27–30]**, NOB **[31–32]**, DNB **[15,33]**. A total of 25 genera and 67 OTUs were found and over 75% of the OTUs were from Proteobacteria.

**Figure 5.**
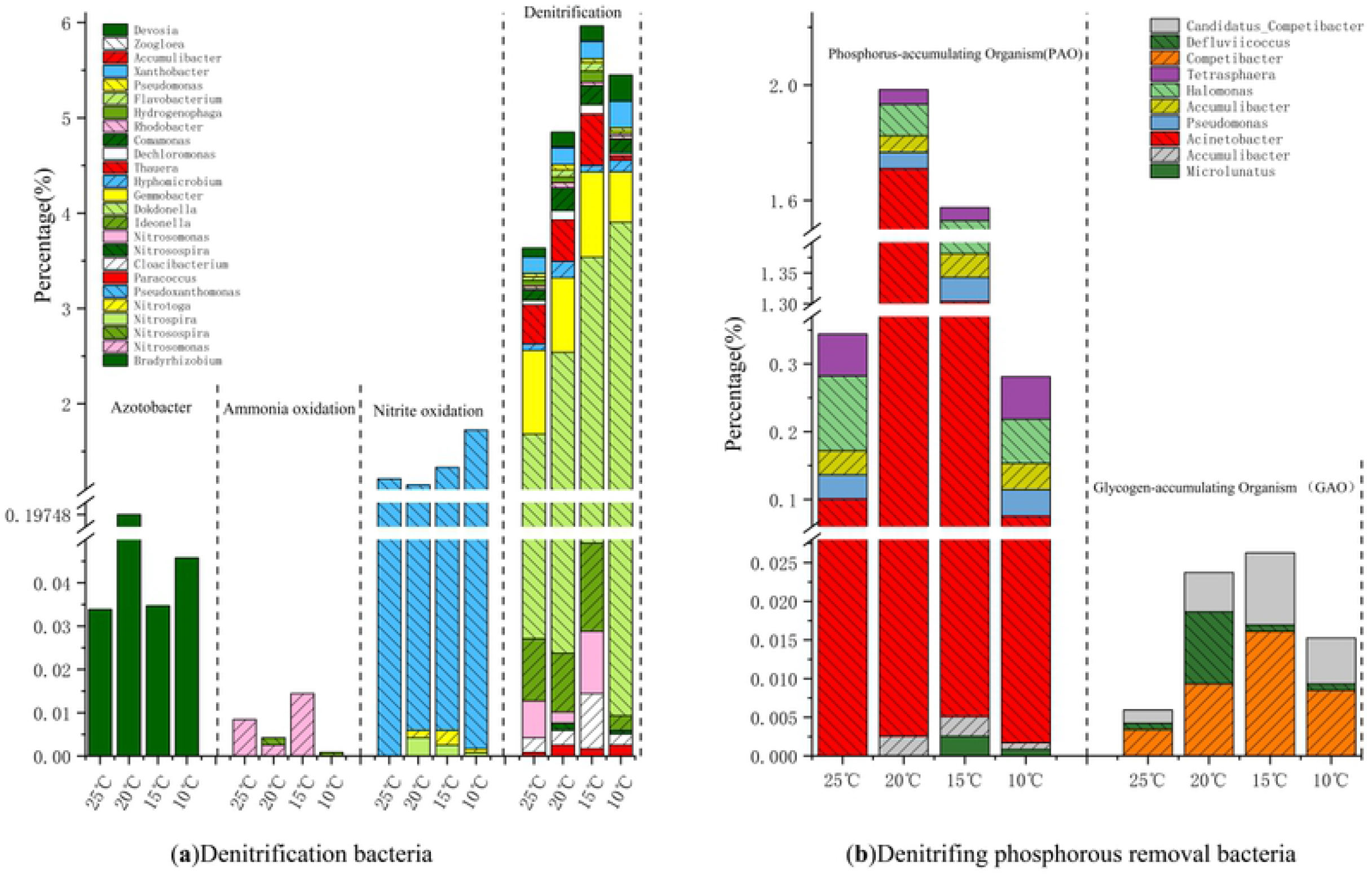
Bacteria on genus level

Azotobacter can transform nitrogen elements in waste water into organic matter to remove nitrogen content in waste water. It has been found that Azotobacter can transform nitrogen elements in waste water into alginate, so as to realize resource recovery of waste water treatment **[34]**. Although the overall content of Azotobacter is low (< 0.2%), the content at 20°C is about 5 times that of other temperatures. 20°C is the temperature condition suitable for Azotobacter bacteria enrichment in the plateau temperature environment.

By oxidizing ammonia nitrogen to nitrite nitrogen, AOB participates in nitrification reaction**[35]**, which is the important step that limits the rate of nitrification in nitrification removal. AOB contains 5 genera **[30]**and in this sequencing, the AOB were composed of Nitrosomonas and Nitrosospira, with the lowest content at 10°C. The removal rate of AOB has been low, which is directly related to the low content of AOB (< 0.02%) in this experiment. The removal rate of total nitrogen and ammonia nitrogen tended to be stable at 15°C and decreased significantly at 10°C. This is consistent with the abundance change of ammonia-oxidizing bacteria.

By oxidizing nitrite nitrogen to nitrate nitrogen, NOB participates in denitrification reaction. Nitrite is harmful to organisms and carcinogenic to humans **[36]**. In water aquaculture industry, it is an important task to treat high-nitrate nitrogen waste water **[37]**, and the abundance of nitrite bacteria is crucial. However, in short-range nitrification, controlling the nitrate nitrogen abundance has become an important task to be considered **[38]**. Pseudoxanthomonas from γ-proteobacteria dominated the Pseudoxanthomonas genus (> 99%) with little change in abundance under four temperature conditions. As reported in many experiments, Pseudoxanthomonas has better degradation ability compared with common non-steroidal anti-inflammatory drugs (NSAIDs), such as Diclofenac (DCF), Ibuprofen (IBU), and Naproxen (NAP) **[39]**, and has higher Biosurfactants production potential **[40]**.

Denitrifying bacteria reduces nitrate to nitrite, NO, N2O and nitrogen in turn **[41]**. In this experiment, a total of 20 genera and 56 OTUs were detected, and the proportions of abundance were 74.5%, 78.9%, 81.3% and 75.6% at 4 temperature conditions, respectively, which were the nitrogen removal functional genera with the most abundance in this sequencing. Generally, denitrifying bacteria showed a rising trend at 25°C and 15°C working conditions and decreasing at 10°C. The abundance of 17 strains decreased significantly at 10°C. The dominant genus, Dokdonella, has been reported to have a significant effect on nitrobenzene removal**[42–43]**. The increment of Dokdonella with the decrease of temperature had a significant influence on the increase of the abundance of denitrifying bacteria at low temperature. The actual detection times of Dokdonella at 25°C, 20°C, 15°C and 10°C were respectively 1465, 2007, 3126, 3257, showing its adaptability to the low temperature environment. However, there is no relevant report about that adaptability, which may be related to its source from natural low temperature waste water.

According to the actual phosphorus removal effect, it was divided into Phosphorous-accumulating Organism(PAO) with actual phosphorus removal effect and Glycogen-accumulating Organism (GAO) without phosphorus removal effect. A total of 10 bacteria genera and 26 OTUs were found(Figure 5 **b**), and over 90% of the OTUs came from Proteobacteria. PAO can absorb volatile fatty acids (VFA) by hydrolyzing phosphorous particles in cells under anaerobic conditions, and synthesize polyhydroxyalkyd acid (PHA) by reducing force generated by glycogen decomposition. Under aerobic conditions, PHA can be used to synthesize glycogen, grow and reproduce by consuming the energy generated by PHA, and absorb excessive phosphorus at the same time.

Finally, phosphorus removal can be achieved through mud discharge **[44]**. The competition between the polysaccharides and the polyphosphate bacteria is formed in the anaerobic environment. Generally, the aggregation of GAO is not conducive to phosphorus removal. However, it has also been reported that the abundance of GAO exceeds that of PAO at 32°C, but the removal effect is not affected **[45]**. Factors known that affect the abundance of PAO and GAO include carbon source, temperature and pH value **[46–48]**. At 25°C, the abundance of both PAO and GAO was low; however, it increased significantly at 20°C and 15°C, but decreased again at 10°C. It has been reported that PAO can achieve better phosphorus removal effect at low temperature **[49]**, which is consistent with the results of waste water quality analysis in this experiment. Acinetobacter was the dominant bacterium in this sequencing. Acinetobacter is a traditional dominant polyphosphorous bacterium. It has been reported that it has a good degradation effect on complex molecules with high phosphorus content, such as fats and oils**[50]**. Some bacteria among Acinetobacter genus have the denitrification effect **[51]**. Both Accumulibacter and Tetrasphaera were the dominant bacteria reported in recent years in the enhanced biological phosphorus removal (EBPR) system, in which the content of Accumulibacter was lower, while the abundance of Tetrasphaera was higher than that of Accumulibacter, which was consistent with the study **[52]**.

## Conclusion

1. Plateau temperature has a small influence on the COD and TP removal rate, but a large influence on that of TN and NH4+-N. TP can achieve stable and high removal effect at low temperature, and the removal effect of TN and NH4+ -n weakens seriously at 15°C and 10°C.
2. Compared with the working conditions of 20°C and 15°C, the removal effect at 25°C and 10°C is not stable, and the homogeneity of microbial species is poor.
3. The rapidly changing temperature has little influence on the overall level abundance on phylum level. The phylum composition is relatively similar under all temperature conditions. Bacteroidetes, Proteobacteria and Actinobacteria were the dominant phyla. The functional bacteria related to nitrogen and phosphorus removal basically come from γ-Proteobacteria under Proteobacteria, and the dominant bacteria have obvious advantages and uneven distribution.
4. The abundance of related bacteria in nitrogen removal first increased and then decreased. NOB and DNB were more abundant than AOB and Azotobacter.
5. The total abundance of PAO and GAO first increased and then decreased, and the overall advantage of PAO was greater than that of GAO under plateau temperature condition.
6. A large number of dominant bacteria are resistant, such as Pseudoxanthomonas, which is highly adaptive to the environment of antibiotics, and Dokdonella, which is more resistant to toxic substances represented by nitrobenzene, which is possibly related to water quality and needs further research.

## Reference

[1] Ritts, B. D., Yue, Y., Graham, S. A., Sobel, E. R., Abbink, O. A., & Stockli, D. From sea level to high elevation in 15 million years: Uplift history of the northern Tibetan Plateau margin in the Altun Shan. J.American Journal of Science 308, 657–678 (2008).

[2] Zong, Y., Li, Y., Hao, K., Lu, G. & Huang, D. INFLUENCE OF TRANSIENT CHANGE OF WATER TEMPERATURE ON PILOT-SCALE ANAEROBIC-ANOXIC-OXIC PROCESS UNDER PLATEAU ENVIRONMENTAL FACTORS. J.APPLIED ECOLOGY AND ENVIRONMENTAL RESEARCH 17, 12191–12202 (2019).

[3] Nguyen, L. N., Nghiem, L. D., Pramanik, B. K. & Oh, S. Cometabolic biotransformation and impacts of the anti-inflammatory drug diclofenac on activated sludge microbial communities. J.Science of the Total Environment 657, 739–745 (2019).

[4] Ju F, Zhang T. Advances in meta-omics research on activated sludge microbial community. J Microbiology China, 2019, 46(08):2038–2052 (2019).

[5] Li, C., Liu, S., Ma, T., Zheng, M. & Ni, J. Simultaneous nitrification, denitrification and phosphorus removal in a sequencing batch reactor (SBR) under low temperature. J.Chemosphere 229, 132–141 (2019).

[6] Zhang, X.-d., Song, Q.-w. & Dai, J.-g. WANG Zhi-hui Engineering Design Center of Chinese Research Academy of Environmental Sciences, Beijing 100012, China; Effect of Sewage Temperature and Carbon Source for Nitrogen and Phosphorus Removal Rate of NPR Process. J. Research of Environmental Sciences 4 (2007).

[7] van der Waals, M. J. et al. Ethyl tert-butyl ether (EtBE) degradation by an algal-bacterial culture obtained from contaminated groundwater. J. Water research 148, 314–323 (2019).

[8] Donghui Li, Weiguang Li, Kailei Zhang, Guanglin Zhang, Houqiang Zhang, Duoying Zhang, Pengfei Lv, Jiao Wu. Nutrient removal by full-scale Bi-Bio-Selector for Nitrogen and Phosphorus removal process treating urban domestic sewage at low C/N ratio and low temperature conditions. J. Process Safety and Environmental Protection (2020).

[9] Talinli, I. & Anderson, G. Interference of hydrogen peroxide on the standard COD test. J.Water research 26, 107–110 (1992).

[10] Yi, F. Study on the Influencing Factors of Total Nitrogen Determination in Water QualityAnalysis. J. Guangzhou Chemistry 3 (2012).

[11] Oon, Y.-L., Ong, S.-A., Ho, L.-N., Wong, Y.-S., Dahalan, F. A. Constructed wetland-microbial fuel cell for azo dyes degradation and energy recovery: Influence of molecular structure, kinetics, mechanisms and degradation pathways. J. Science of The Total Environment 720, 137370 (2020).

[12] Qunhua, H. Advances on the analysis methods of ammonia nitrogen in water. J.Guangdong Chemical Industry 14 (2013).

[13] Dai, Z., Zhu, H. & Wen, F. Two nonparametric approaches to mean absolute deviation portfolio selection model. J. Journal of Industrial & Management Optimization 13, 1 (2019).

[14] Zhou, H. & Xu, G. Integrated effects of temperature and COD/N on an up-flow anaerobic filter-biological aerated filter: Performance, biofilm characteristics and microbial community. J.Bioresource technology 293, 122004 (2019).

[15] State Environmental Protection Administration ed. Discharge standard of pollutants for municipal wastewater treatment plant(GB 18918 – 2002). Beijing: China Environmental Science Press, 2002.

[16] Aït-Sahalia, Y. & Xiu, D. Principal component analysis of high-frequency data. J. Journal of the American Statistical Association 114, 287–303 (2019).

[17] Nielsen, P. H., McIlroy, S. J., Albertsen, M. & Nierychlo, M. Re-evaluating the microbiology of the enhanced biological phosphorus removal process. J. Current opinion in biotechnology 57, 111–118 (2019).

[18] Haosagul, S., Prommeenate, P., Hobbs, G. & Pisutpaisal, N. Sulfide-oxidizing bacteria community in full-scale bioscrubber treating H2S in biogas from swine anaerobic digester. J.Renewable Energy 150, 973–980 (2020).

[19] Zhong, C., Li, J., Flynn, S. L., Nesbø, C. L., Sun, C., von Gunten, K.,… Alessi, D. S. Temporal Changes in Microbial Community Composition and Geochemistry in Flowback and Produced Water from the Duvernay Formation. J.ACS Earth and Space Chemistry 3, 1047–1057 (2019).

[20] Zhang, G., Su, F., Jiao, Y., Chen, Q. & Lee, D.-J. Biocathodic performance of bioelectrochemical systems operated at low temperature. J.Bioresource Technology, 123463 (2020).

[21] Nathani, N. M., Mootapally, C. & Dave, B. P. Antibiotic resistance genes allied to the pelagic sediment microbiome in the Gulf of Khambhat and Arabian Sea. J.Science of The Total Environment 653, 446–454 (2019).

[22] Gu, Y., Wei, Y., Xiang, Q., Zhao, K., Yu, X., Zhang, X.,… Zhang, X. C: N ratio shaped both taxonomic and functional structure of microbial communities in livestock and poultry breeding wastewater treatment reactor. J. Science of the Total Environment 651, 625–633 (2019).

[23] Liu, J., Wang, C., Wu, K., Huang, L., Tang, Z., Zhang, C.,… Yang, B. Novel start-up process for the efficient degradation of high COD wastewater with up-flow anaerobic sludge blanket technology and a modified internal circulation reactor. J.Bioresource Technology, 123300 (2020).

[24] Ghosh, U. D., Singh, P. K., Ganguli, S., Saha, C., Chandra, A., Seal, A., & Ghosh, M. M. Rhizospheric soil of Typha angustifolia L. from heavy metal contaminated and free sites. J.Comparative profiling reveals selective abundance of γ-proteobacteria and β-proteobacteria. (2019).

[25] Zheng, M., Wang, M., Zhao, Z., Zhou, N., He, S., Liu, S.,… Wang, X. Transcriptional activity and diversity of comammox bacteria as a previously overlooked ammonia oxidizing prokaryote in full-scale wastewater treatment plants. J.Science of the Total Environment 656, 717–722 (2019).

[26] Huang, J.-f., Zhang, D.-f., Leng, B., Lin, Z.-c. & Pan, Y.-t. Response surface optimization of conditions for culturing Azotobacter chroococcum in Agaricus bisporus industrial wastewater. J.The Journal of general and applied microbiology 65, 163–172 (2019).

[27] Ju, F., Xia, Y., Guo, F., Wang, Z. & Zhang, T. Taxonomic relatedness shapes bacterial assembly in activated sludge of globally distributed wastewater treatment plants. J.Environmental microbiology 16, 2421–2432 (2014).

[28] Juretschko, S., Timmermann, G., Schmid, M., Schleifer, K.-H., Pommerening-Röser, A., Koops, H.-P., & Wagner, M. Combined molecular and conventional analyses of nitrifying bacterium diversity in activated sludge: Nitrosococcus mobilis and Nitrospira-like bacteria as dominant populations. J. Appl. Environ. Microbiol. 64, 3042–3051 (1998).

[29] McIlroy, S. J. et al. MiDAS: the field guide to the microbes of activated sludge. J.Database 2015 (2015).

[30] Koops, H. P. & Pommerening-Röser, A. The Lithoautotrophic Ammonia-Oxidizing Bacteria. J.Prokaryotes, 1–17 (2015).

[31] Saunders, A. M., Albertsen, M., Vollertsen, J. & Nielsen, P. H. The activated sludge ecosystem contains a core community of abundant organisms. J.The ISME journal 10, 11 (2016).

[32] Nowka, B., Daims, H. & Spieck, E. Comparison of oxidation kinetics of nitrite-oxidizing bacteria: nitrite availability as a key factor in niche differentiation. J.Appl. Environ. Microbiol. 81, 745–753 (2015).

[33] Thomsen, T. R., Kong, Y. & Nielsen, P. H. r. Ecophysiology of abundant denitrifying bacteria in activated sludge.J. FEMS microbiology ecology 60, 370–382 (2007).

[34] Aarstad, O. A., Stanisci, A., SÃ¦trom, G. I., TÃ.ndervik, A., Sletta, H. v., Aachmann, F. L., & SkjÃ¥k-BrÃ¦k, G. Biosynthesis and Function of Long Guluronic Acid-blocks in Alginate produced by Azotobacter vinelandii. J.Biomacromolecules, 20(4), 1613–1622 (2019).

[35] Zheng, M., Wang, M., Zhao, Z., Zhou, N., He, S., Liu, S.,… Wang, X. Transcriptional activity and diversity of comammox bacteria as a previously overlooked ammonia oxidizing prokaryote in full-scale wastewater treatment plants. J.Science of the Total Environment, 656, 717–722 (2019).

[36] Alonso, H. R. F. et al. USING SALIVARY NITRITE LEVELS AS A POTENTIAL BIOMARKER FOR ORAL CHEMICAL CARCINOGENESIS. J.Oral Surgery, Oral Medicine, Oral Pathology and Oral Radiology 129, e180 (2020).

[37] Holmes, D. E., Dang, Y. & Smith, J. A.J. Nitrogen cycling during wastewater treatment. J.Advances in applied microbiology Vol. 106 113–192 (Elsevier, 2019).

[38] Shi, W., Li, H. & Li, A. Mechanism and influencing factors of nitrogen removal in subsurface flow constructed wetland. J.Applied Chemical Engineering 1 (2018).

[39] Lu, Z., Sun, W., Li, C., Ao, X., Yang, C., & Li, S. Bioremoval of non-steroidal anti-inflammatory drugs by Pseudoxanthomonas sp. DIN-3 isolated from biological activated carbon process. J.Water research, 161, 459–472 (2019).

[40] Astuti, D. I., Purwasena, I. A., Putri, R. E., Amaniyah, M. & Sugai, Y. Screening and characterization of biosurfactant produced by Pseudoxanthomonas sp. G3 and its applicability for enhanced oil recovery. J. Journal of Petroleum Exploration and Production Technology 9, 2279–2289 (2019).

[41] Zhang, H., Feng, J., Chen, S., Zhao, Z., Li, B., Wang, Y.,… Yan, M. Geographical patterns of nirS gene abundance and nirS-type denitrifying bacterial community associated with activated sludge from different wastewater treatment plants. J.Microbial ecology, 77(2), 304–316 (2019).

[42] Jiang, X., Shen, J., Xu, K., Chen, D., Mu, Y., Sun, X.,… Wang, L. Substantial enhancement of anaerobic pyridine bio-mineralization by electrical stimulation. J.Water research, 130, 291–299 (2018).

[43] Liang, B., Qi, M., Yun, H., Zhao, Y., Bai, Y., Kong, D., & Wang, A.-J. Electrode-Respiring Microbiomes Associated with the Enhanced Bioelectrodegradation Function. J.Bioelectrochemistry Stimulated Environmental Remediation (pp. 47–72): Springer (2019).

[44] Mino, T., Van Loosdrecht, M. & Heijnen, J. Microbiology and biochemistry of the enhanced biological phosphate removal process. J.Water research 32, 3193–3207 (1998).

[45] Ong, Y. H., Chua, A. S. M., Fukushima, T., Ngoh, G. C., Shoji, T., & Michinaka, A. High-temperature EBPR process: The performance, analysis of PAOs and GAOs and the fine-scale population study of Candidatus â€œAccumulibacter phosphatisâ€□. J.Water research 64, 102–112 (2014).

[46] Lopez-Vazquez, C. M., Oehmen, A., Hooijmans, C. M., Brdjanovic, D., Gijzen, H. J., Yuan, Z., & van Loosdrecht, M. C. Modeling the PAO–GAO competition: effects of carbon source, pH and temperature.J. Water Research, 43(2), 450–462 (2009).

[47] Carvalheira, M. n., Oehmen, A., Carvalho, G. & Reis, M. A. Survival strategies of polyphosphate accumulating organisms and glycogen accumulating organisms under conditions of low organic loading. J.Bioresource technology 172, 290–296 (2014).

[48] Smolders, G., Van der Meij, J., Van Loosdrecht, M. & Heijnen, J. Model of the anaerobic metabolism of the biological phosphorus removal process: stoichiometry and pH influence. J.Biotechnology and bioengineering 43, 461–470 (1994).

[49] Tian, W.-D., Lopez-Vazquez, C., Li, W.-G., Brdjanovic, D. & Van Loosdrecht, M. Occurrence of PAOI in a low temperature EBPR system. J.Chemosphere 92, 1314–1320 (2013).

[50] Jyoti, R., Ramesh, C., Chandra, S. D., Supriya, T., Saurabh, K., & Reeta, G. Comparative phosphate solubilizing efficiency of psychrotolerant Pseudomonas jesenii MP1 and Acinetobacter sp. ST02 against chickpea for sustainable hill agriculture. J.Biologia (2018).

[51] Yang, J., Wang, Y., Chen, H. & Lyu, Y. Ammonium removal characteristics of an acid-resistant bacterium Acinetobacter sp. JR1 from pharmaceutical wastewater capable of heterotrophic nitrification-aerobic denitrification. J.Bioresource Technology 274, 56–64 (2019).

[52] Liu, R., Hao, X., Chen, Q. & Li, J. Research advances of Tetrasphaera in enhanced biological phosphorus removal: A review. J.Water Research 166, 115003 (2019).

